# Visual plasticity and exercise revisited: no evidence for a “cycling lane”

**DOI:** 10.1101/448498

**Authors:** Abigail E Finn, Alex S Baldwin, Alexandre Reynaud, Robert F Hess

**Affiliations:** McGill Vision Research Department of Ophthalmology McGill University Montreal, Quebec, Canada

**Keywords:** vision, psychophysics, binocular, rivalry, exercise, ocular dominance, plasticity, monocular deprivation

## Abstract

Experiments using enriched environments have shown that physical exercise modulates visual plasticity in rodents. A recent study (Lunghi & Sale, 2015, doi: 10.1016/j.cub.2015.10.026) investigated whether exercise also affects visual plasticity in adult humans. The plastic effect they measured was the shift in ocular dominance caused by 2 hours of monocular deprivation (e.g. by an eye patch). They used a binocular rivalry task to measure this shift. They found that the magnitude of the shift was increased by exercise during the deprivation period. This effect of exercise was later disputed by a study that used a different behavioural task (Zhou *et al.*, 2017, doi: 10.1155/2017/4780876). Our goal was to determine whether the difference in task was responsible for that study’s failure to find an exercise effect. We set out to replicate Lunghi & Sale (2015). We measured ocular dominance with a rivalry task before and after 2 hours of deprivation. We measured data from two conditions in 30 subjects. On two separate days they either performed exercise or rested during the deprivation period. Contrary to the previous study, we find no significant effect of exercise. We hypothesise that exercise may affect rivalry dynamics in a way that interacts with the measurement of the deprivation effect.

## 1 Introduction

Brain plasticity is usually thought of in terms of critical periods in young animals. It is now recognized, however, that a degree of plasticity remains into adulthood. Studies using rodents have shown experience-dependent plasticity as a result of enriched environments. These can have both structural and functional benefits for the adult brain (Sale et al., 2016). One key component of these enriched environments is physical exercise. Previous studies have also found that exercise increases neural responsivity in the visual cortex of adult rodents (Niell and Stryker, 2010).

While exercise effects have been found in the visual cortex of rodents (Sale et al., 2007), their extrapolation to primates is not conclusive. In humans, evidence for an exercise effect on plasticity has previously been limited to prefrontal and hippocampal regions (Erickson et al., 2013). This was the case until the recent study by Lunghi and Sale (2015). They argued that exercise could modulate the plasticity of early visual cortex of adults. For their measure of plasticity, Lunghi and Sale (2015) used the shift of ocular dominance caused by short-term monocular deprivation. If one eye is deprived of visual input for a period of around two hours, there is a shift in dominance in favour of the patched eye. This effect has been demonstrated with deprivation from eye patches (Lunghi et al., 2011), dichoptically-presented movies (Zhou et al., 2014; Bai et al., 2017), and continuous flash suppression (Kim et al., 2017). The shift in ocular dominance has been confirmed with EEG (Zhou et al., 2015), fMRI (Binda et al., 2018), MEG (Chadnova et al., 2017), and various psychophysical tasks (Zhou et al., 2013; Spiegel et al., 2017; Baldwin and Hess, 2018). For their study on the effect of exercise, Lunghi and Sale (2015) measured the ocular dominance shift with a binocular rivalry paradigm. The perceptual dominance of each eye was monitored before and after deprivation. They found that exercise during the deprivation period enhanced the shift in ocular dominance. However, a later study by Zhou et al. (2017) failed to find an effect of exercise. Their methods were similar to Lunghi and Sale (2015), although rather than binocular rivalry they used a task that measured the relative strength of the input from each eye to a fused binocular percept.

It is possible that the use of a fusion rather than a rivalry task is responsible for the failure to replicate the previous effect. Recent studies have demonstrated that different effects of monocular deprivation may be found depending on the psychophysical task used to measure them (Bai et al., 2017; Baldwin and Hess, 2018). The binocular fusion task used by Zhou et al. (2017) involves two grating stimuli of the same orientation and spatial frequency, but slightly different spatial phase. These are fused into a single binocular percept, with the phase of that perceived grating shifted to favour the dominant eye. This combination can be attributed to excitatory interactions. The rivalry task used by Lunghi and Sale (2015) is the perceptual alternation of stimuli of orthogonal orientation that stimulate different neuronal populations. Their perceptual competition is subserved by inhibitory interactions (Klink et al., 2010). Zhou et al. (2017) left open the possibility that differences between the results from the two studies may be due to the different methods used. One task measures inhibitory interactions between different neural populations tagging left and right eye responses. The other measures excitatory combination of left/right eye signals within a single neuron population.

To determine whether the difference in task was responsible for the previous failure to replicate, we set out to re-assess the effect of exercise on visual plasticity. As before, we used the short-term monocular deprivation paradigm, but with the binocular rivalry measure rather than the binocular fusion measure. This amounts to a replication of the study of Lunghi and Sale (2015). We have endeavored to use as similar a protocol to theirs as their described methods allow.

## 2 Methods

### 2.1 Subjects

We recruited 30 subjects including one author (AF), 2 experienced psychophysicists, and 27 naïve subjects. The average age was 22 years (range 18-26). Of the subjects recruited 21 were female. All procedures were approved by the research ethics board of the McGill University Health Centre, and carried out in accordance with the Declaration of Helsinki. Informed written consent was obtained from the subjects.

### 2.2 Procedures

In the binocular rivalry experiment, oblique Gabor patches at +45 and −45 degrees were presented separately to the two eyes using shutter glasses. The Gabors had a spatial frequency of 1.5 c/deg and a spatial sigma of 1.3 degrees of visual angle and a contrast of 50%. The two grating orientations were randomly assigned to the two eyes for each block to avoid any orientation bias. For a three-minute testing period, subjects then used a keypad to continuously indicate their current percept. Subjects had three buttons to press to signal three perceptual states: “left oblique grating”, “right oblique grating”, and “mixed percept”. All three states were illustrated for the subject before they began testing so they knew how to respond. Data were processed by assigning the two Gabor orientations to the role of the eye in which they were presented (patched or non-patched eye).

We measured the shift in ocular dominance after patching in two conditions. In the control condition subjects rested during the patching period. In the cycling condition subjects performed physical exercise by cycling on a stationary standing bike during the patching period. Subjects performed the two conditions on different days (separated by at least 24 hours). For both conditions, a baseline measurement was made using the binocular rivalry task. After this measurement, a translucent patch was fitted over the dominant eye of the subjects (determined using the Miles test). For 2 hours, the subject then viewed movies (cinematic films) on the projector screen. In the control condition the subject sat in a chair throughout this period. In the cycling condition subjects used a standing bike to alternate 10 minute periods of exercise (target heart rate of 120 beats per minute) and rest. After 2 hours, the patch was removed and subjects were tested at 0, 4, 8, 12, 30, 45, 60, 90, and 120 minutes following patch removal.

### 2.3 Apparatus

Subjects sat at 2.3 m from a projector screen. Both the stimuli for the binocular rivalry task and the movies viewed during the monocular patching period were presented on the screen by an Optoma HD26 DLP projector. The resolution of the projector gave 75 pixels per degree of visual angle. The mean luminance of the screen was 95 cd/m^2^. The experiment was programmed in Matlab using Psychtoolbox (Kleiner et al., 2007). During the binocular rivalry task, the grating stimuli were presented dichoptically using frame-interleaving with a pair of Optoma ZD302 DLP Link Active Shutter 3D Glasses. During the binocular rivalry task, the control movie viewing, and the rest periods in the cycling movie viewing subjects sat on a chair. During the active periods in the cycling condition subjects pedalled a stationary bike at such a pace as to sustain a heart rate of 120 beats per minute (monitored by a Polar H7 Heart Rate Sensor).

## 3 Results

### 3.1 Our replication attempt

Measurements taken over the 120 minutes of recovery from two hours of patching are presented in Figure 1A. The response duration histograms were heavily skewed (Figure 2A-B). To give a measure that is independent of the shape of the histogram, we used the total time that the subject responded that they were seeing the stimulus for the patched or non-patched eye. We used this to calculate the ocular dominance index after Dieter, Sy & Blake (Dieter et al. (2017); Equation 1)

**Figure 1.**
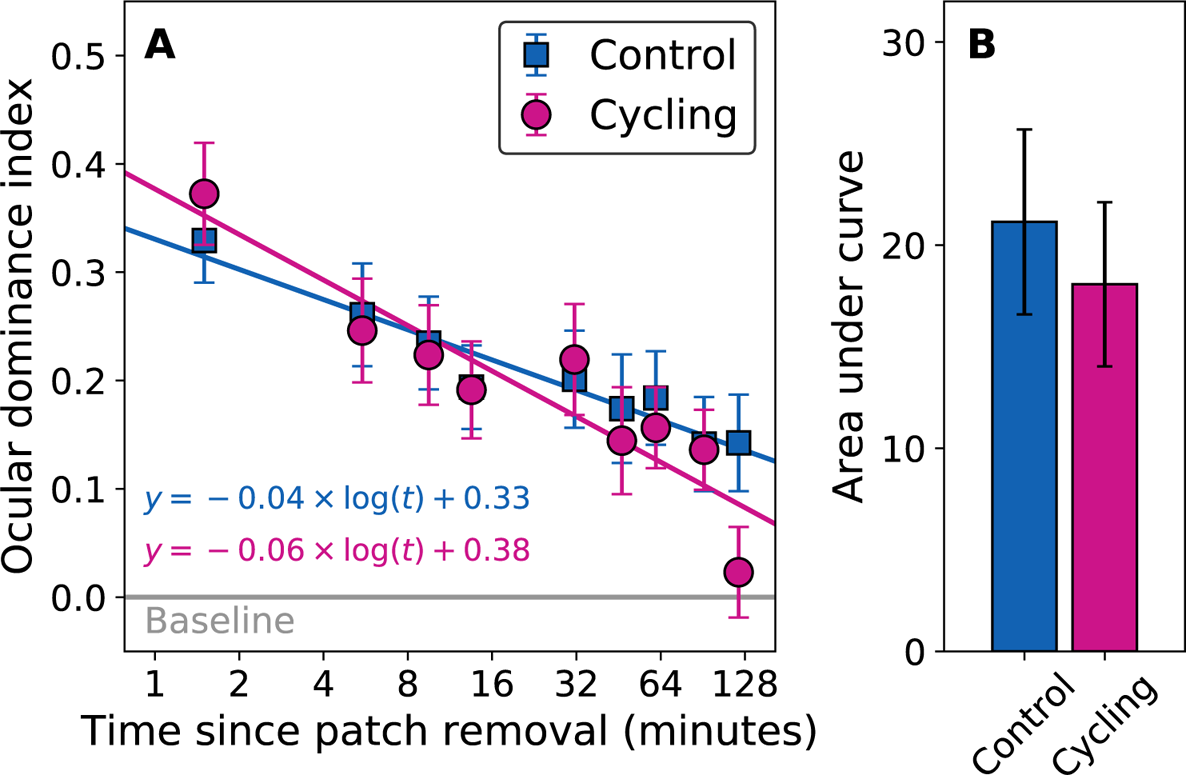
Panel A shows the time-course of recovery from patching (each time-point is given as the middle of the 3 minute measurement period). The ocular dominance index (ODI) is calculated from the total times for which the patched and non-patched eye percepts were seen. For each subject, the ODIs after patching were normalised by the baseline before patching. An ODI of zero means no change. Increases favour the patched eye. The mean over 30 subjects is plotted (error bars show standard error). Data were fit by a straight line on semi-log axes, with equations provided in the figure. Panel B shows the overall effect of patching (mean +/− standard error). This is the trapezoidal area between the data points for each subject and their baseline on a linear time axis.

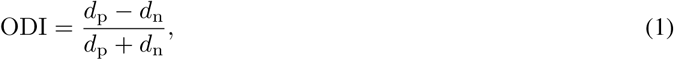

where *d*_p_ and *d*_n_ are the total response durations for the patched and non-patched eyes respectively.

The time-courses of the recovery from 2 hours of patching with and without exercise are presented in Figure 1A for the ODI calculation. Five of our thirty subjects each had a single datapoint where the data were not collected. Where a subject was missing a datapoint for one condition we also excluded data from the same timepoint in the other condition. These five subjects were also excluded from the two-way repeated-measures ANOVA analysis we performed in RStudio (RStudio Team, 201 6). The results from the ANOVA show a significant effect of time-point (F(8,192) = 11.09, p < 0.001) but no effect of exercise (F(1,24) = 0.82, p = 0.37), and no interaction effect (F(8,192) = 1.19, p = 0.31).

We performed an additional analysis taking the area under the ratio vs. linear time curve (Figure 1B). The area gives the overall deviation from baseline, representing the magnitude of the patching effect. We calculated the area for each subject (normalised for their individual baseline measurements). As removal of a single timepoint did not prevent an area being calculated, all subjects’ data were used in this analysis. We found no significant difference between the cycling and control conditions (t(29) = −0.84, p = 0.41). In fact, in our data the (non-significant) trend is in the opposite direction, with a reduced patching effect in the cycling condition.

### 3.2 Comparison with Lunghi & Sale

In addition to the ocular dominance index, we considered two other ways to analyse the binocular rivalry data. In Lunghi and Sale (2015) the mean durations of the individual “patched eye” and “non-patched eye” percepts were used to calculate the ocular dominance ratios. The duration distributions were skewed however (Figure 2A-B), so the mean would be affected by the relatively few responses that have much longer durations than the others. To account for this, we also analysed the median response duration. To calculate the ocular dominance, we divided the average (mean or median) response duration for the patched eye by the average duration for the non-patched eye. We took the log_2_ of that ratio, so that a 2:1 ratio in favour of the patched eye would be a log_2_ ratio of 1, whereas a 1:2 ratio in favour of the non-patched eye would give a log_2_ ratio of −1. For the mean and median ratio measures, the recovery from patching was similar to Figure 1A and gave the same pattern of results on the ANOVA analysis.

**Figure 2.**
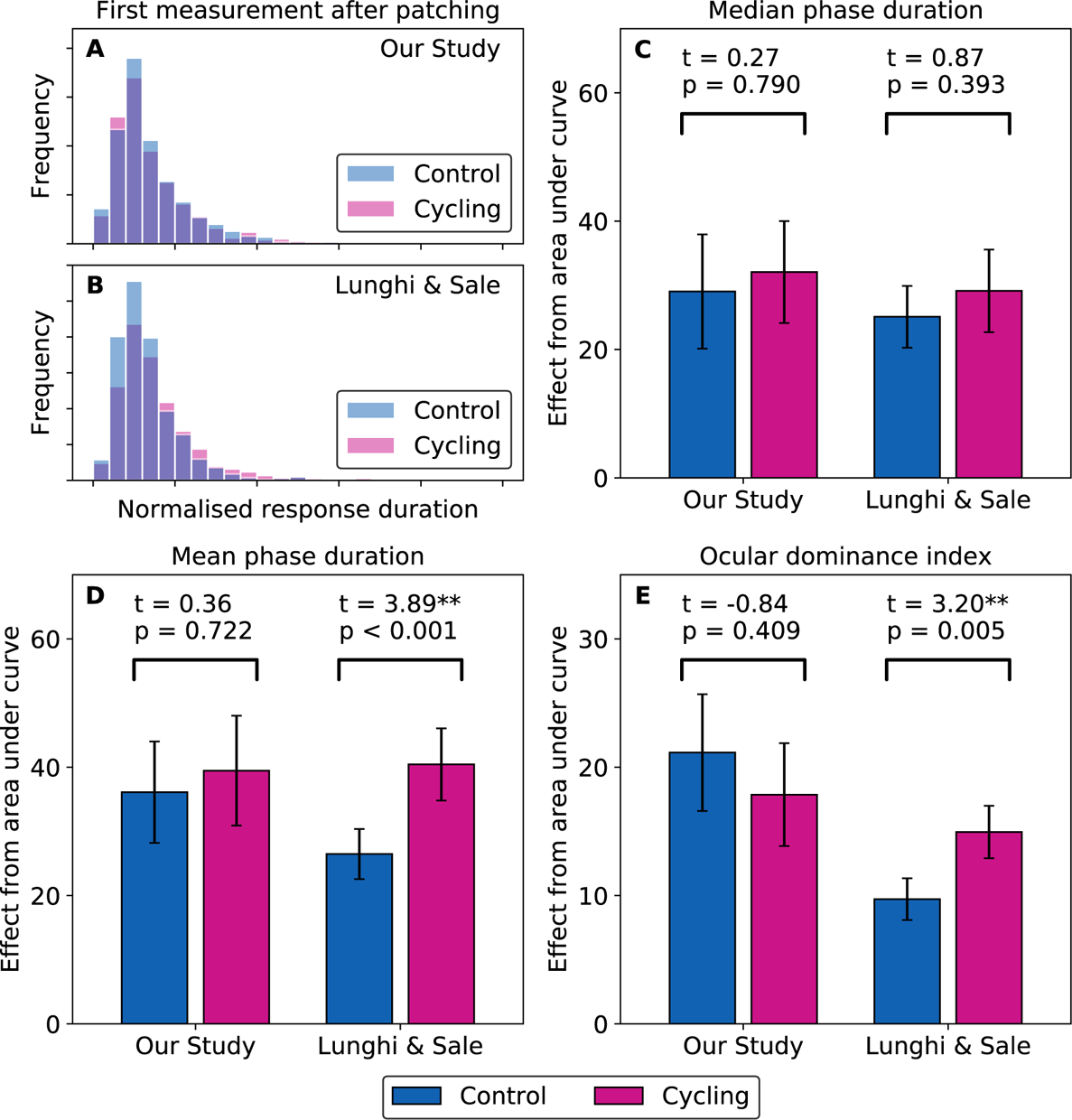
Panels A-B show histograms of patched eye response durations following patching (pooled across all subjects). In Panel A, the data are from measurements made immediately after patch removal in our study. These durations are normalised by the median duration for the patched eye in the baseline data. In Panel B the data used are from Lunghi and Sale (2015), taken at the first timepoint after patch removal (8 minutes). Histograms from the cycling and control conditions are shown overlaid transparently on top of each other. Panels C-E show mean-across-subjects patching effects in the control and cycling conditions (error bars show standard error). These are calculated as the area under the “eye dominance” vs. time curves. Three measures are used to calculate eye dominance. For each measure, we have compared the overall cycling vs. control effect with a paired t-test. In C, the eye dominance is calculated as the ratio between the median response durations for patched and non-patched eye percepts. In D, the same calculation is made with the mean response durations. In E, we calculated the ocular dominance index from the total time during which the patched and non-patched eye percepts were seen.

In Figure 2C-E we present the mean-across-subjects values for the median duration, mean duration, and ocular dominance index measures. For each study, we performed a paired t-test in SciPy (Jones et al., 2001). In our data, we find no significant difference between the cycling and control conditions. In the data from Lunghi and Sale (2015) we find highly significant effects when the mean and ocular dominance index are used as the measures of eye dominance, but no significant effect with the median response duration. These t-tests were used as the basis for a replication Bayes factor calculation (Verhagen and Wagenmakers, 2014) in RStudio (RStudio Team, 2016; Harms, 2018). We used the t-statistics for the ocular dominance index measure to quantify the evidence for the Lunghi and Sale (2015) effect in our data, relative to the null hypothesis where exercise does not affect the patching effect. The ratio between the alternative and null hypotheses was 0.015, giving strong (Kass and Raftery, 1995) support to the null.

The use of these multiple measures allows us to draw a more nuanced conclusion about both the nature of the patching effect and its modulation by exercise. In both studies patching increases the duration of the median patched eye response relative to the median non-patched eye response. It also increases the total proportion of time during which patched eye responses are given. We were not able to find any significant effect of exercise in our data for any of the three measures we analysed. Even the trend favouring a stronger effect in the cycling condition that we find for the median and mean analyses is reversed when we make our ocular dominance index analysis. In the Lunghi and Sale (2015) data there is no significant effect on the median response duration from exercise. This means that there is no evidence that exercise increases the average response duration seen by the patched eye. Instead, it may be that exercise results in an increased duration of the longer responses.

### 3.3 Analysis of the baseline behaviour of the subjects

In our experiment, data were collected from the thirty subjects before analysis was conducted. We did not continuously monitor for any aberrant results or indications of poor data quality. It is possible that had we done so our subject pool would have been reduced to one where the effect of exercise would be found. One possible metric that could have been considered for this purpose would be the proportion of “mixed” percepts reported by a subject. When more mixed percepts are reported, there will be less data regarding the dominance of the “left eye” or “right eye” percepts. These subjects may therefore show a weaker effect of patching and dilute the result. We performed a further analysis of our data to determine whether mixed percepts were responsible for our failure to replicate the results of Lunghi and Sale (2015). We calculated the mixed percepts index (MPI) as

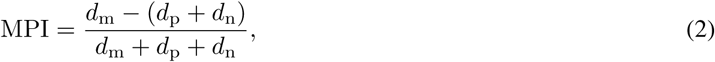

where *d*_p_ and *d*_n_ are the total response durations for the patched and non-patched eyes, and *d*_m_ is the total response duration for the mixed percepts. The median MPI calculated across subjects is presented in Figure 3A. The horizontal black line shows the median MPI from the Lunghi and Sale (2015) data, whereas the rightmost grey circle show the median MPI from our data (30 subjects). As the grey curve declines to the left of that point, it shows the effect of excluding subjects who showed the greater proportion of mixed percepts from our analysis. It is worth noting that only one of our subjects shows a proportion of mixed percepts lower than the *median* subject from Lunghi and Sale (2015). The proportion of mixed percepts will be largely dependent on the criterion that the subject uses. A previous study instructed subjects to only identify the dominant grating when it was *exclusively* visible. They found that their subjects always reported mixed percepts for at least one-third of the duration (O’Shea et al., 1997). Plotting this on Figure 3A would give a horizontal line at −0.33. Our proportion of mixed percepts was lower, as we instructed our subjects to respond based on whichever grating was dominant. They were only to give the “mixed” response when the two appeared to be equally dominant. If similar instructions were given in Lunghi and Sale (2015), then the differences between our studies are consistent with our subjects having different criterion levels for this “mixed” percept. Figure 3C shows how our conclusions regarding the effect of exercise on the ocular dominance shift from patching may have changed if we had excluded subjects who reported a greater proportion of mixed percepts. Only by excluding most of our subjects (more than two-thirds) do we see a non-significant trend for exercise to increase the shift in ocular dominance. However, we would have no principled reason for setting the exclusion criterion at this point.

**Figure 3.**
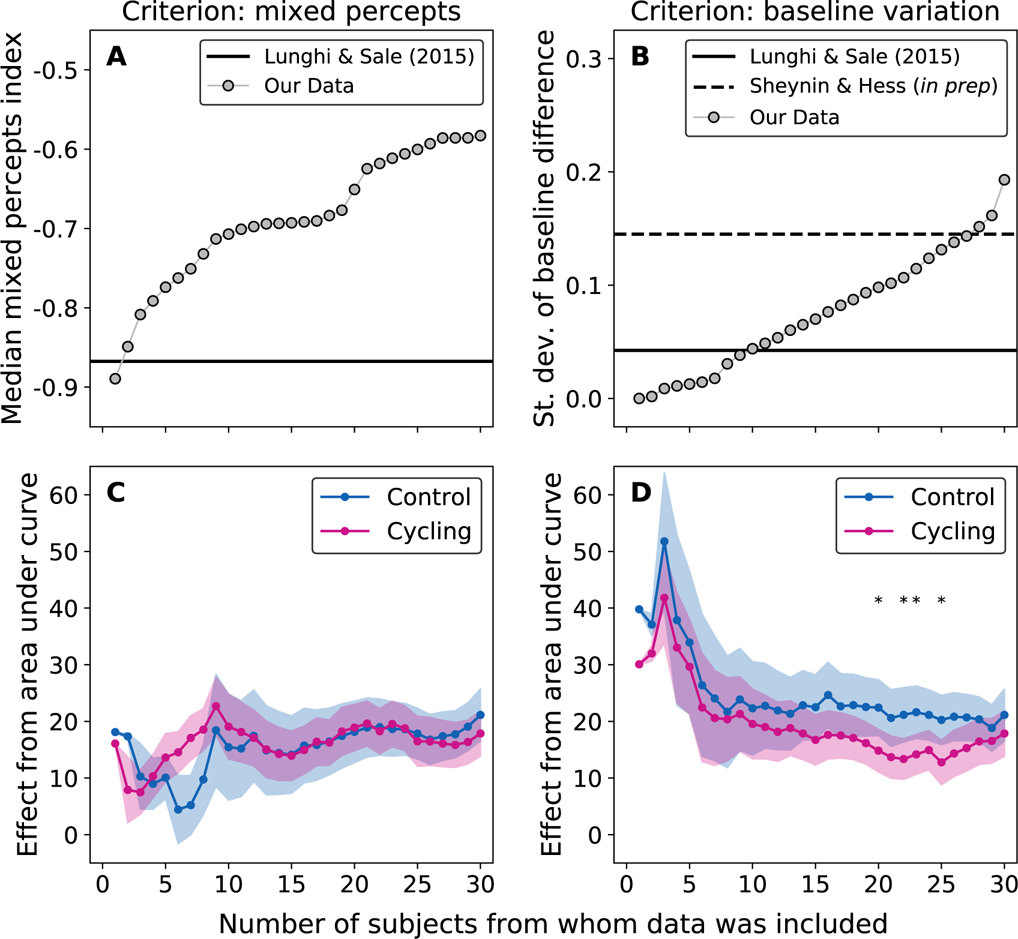
The top two panels present statistics from the baseline data collected in our study, compared to that from Lunghi & Sale Lunghi and Sale (2015). Panel A shows the median mixed percepts index, and how this reduces in our data as we remove subjects with larger proportions of “mixed percept” responses. Panel B shows the standard deviation of the differences between the baseline ODIs measured in the cycling and control sessions. The effect of excluding subjects showing greater day-to-day variation is shown for our data. We also show an equivalent standard deviation calculated from another study (Sheynin and Hess, in prep). In the bottom two panels, the overall patching effect in the control and cycling conditions is given as the area under the curve from the ocular dominance index analysis. The circles show the mean across subjects for each n, with the shaded region giving the standard error. In panel C, the effect of excluding subjects with a greater proportion of “mixed percept” responses (A) is demonstrated. For each group of subjects considered in panel A, the results based on analysing only their data are given by the two points at the same x-axis location in Panel C. Panel D shows the effect of excluding subjects with the greatest shift in baseline dominance ratio between days (B). Points which are significantly different by a paired t-test are marked with asterisks.

Another metric would be the day-to-day variation in the measured baseline. A large difference in baseline between sessions may indicate that data measured from that subject are not reliable. This could add noise to the results, which would also reduce any measure of statistical significance. We calculated the day-to-day baseline variation as the difference in the ocular dominance index (ODI) measured on the two days of testing (one day for the cycling condition and one day for the control). The overall spread was characterised by the standard deviation of the differences. This is shown for our study in Figure 3B by the rightmost grey circle (30 subjects), in comparison to the solid black line from Lunghi & Sale’s Lunghi and Sale (2015) data. Similar to Figure 3A, the data points to the left of that rightmost point indicate the effects of removing subjects showing the greater amount of day-to-day variation in their baselines. Our subjects were more variable than those from Lunghi and Sale (2015). To reduce the standard deviation of our data to match theirs we had to exclude two-thirds of our subjects. Some of this variability will come from measurement error, with the rest due to day-to-day variability in ocular dominance. We obtained an estimate of this day-to-day variability using data from another study where 19 subjects were tested for six rivalry sessions (80 seconds each) across two days (Sheynin and Hess, *in prep*). The median ODI across six sessions will have a much smaller measurement error than the ODI from a single session. We therefore used the median ODI from those six sessions to calculate the standard deviation of the day to day differences in ocular dominance. This is presented as the horizontal dashed line on Figure 3B, and is similar to the standard deviation found with our full data set. The standard deviation found by Lunghi and Sale (2015) was much lower. This suggests that the subjects selected in that study had a particularly stable ocular dominance baseline. Panel D shows how our results would have changed if we excluded subjects who showed greater day-to-day baseline variation. There is no evidence that the more variable subjects were hiding any underlying enhancement of the ocular dominance shift. If anything, we find a significant effect in the *opposite* direction for certain combinations of subjects (indicated by asterisks).

## 4 Discussion

We find no effect of exercise on the transient neuroplastic changes caused by monocular deprivation. This result is surprising considering the size of the effect reported by Lunghi and Sale (2015). Their ANOVA found a large effect of exercise (F(1,19) = 9.58, p = 0.006). It is therefore unlikely that the difference between our studies is due to chance. We calculated a replication Bayes factor (Verhagen and Wagenmakers, 2014) using the ReplicationBF package in RStudio (RStudio Team, 2016; Harms, 2018). We found a Bayes factor of 0.015, indicating strong support (Kass and Raftery, 1995) for the null hypothesis (where exercise does not influence the patching effect). This evidence adds to the lack of an exercise effect found in a previous study using a binocular combination method (Zhou et al., 2017), and as-yet unpublished work we have conducted with dichoptic surround suppression.

The procedures and subject population were similar between our study and Lunghi and Sale (2015). Our binocular rivalry statistics were within the normal range (O’Shea et al., 1997). The stimuli had small differences. The gratings we presented were around 30% larger. Although the contrast was the same (50%), the luminance of our display was 2.5 times higher. None of these differences seem likely to account for the absence of an exercise effect. We performed further analyses to see whether an effect exists in a subset of our cohort. We found that our failure to replicate the previous result was not due to including subjects with more day-to-day variability. Nor was it due including subjects who gave more “mixed percept” responses.

We chose the ocular dominance index to present in Figure 1, but an analysis of the mean phase duration did not find a significant effect in our data either. The ocular dominance index analysis was significant for the data of Lunghi and Sale (2015), though a median phase duration analysis was not. The general effect of patching however is seen across all three measures, including median phase duration. Therefore, if exercise were to enhance that effect, we would expect it to have an equal impact on all three measures. It is possible that exercise may further skew the phase duration distributions in binocular rivalry. This would result in an effect that is driven by a small number of extended percepts.

In the light of our results and the additional analysis of the data from Lunghi and Sale (2015), we propose that rather than exercise enhancing the patching effect per se (as they purport), exercise may be affecting the behaviour of the subjects in a binocular rivalry task in a way that inflates some measures of the patching effect. Controlling for this would require a study that investigated the effect exercise has on binocular rivalry dynamics in the absence of patching.

## 5 Acknowledgements

This work was funded by an ERA-NET Neuron grant (JTC 2015), and CIHR grants (CCI-125686 and 228103) awarded to Robert F. Hess. We would like to thank Claudia Lunghi and Alessandro Sale for sharing the data from their study, and for helpful discussions.

## 6 Author Contributions

This study was designed and conceived by A. E. Finn, A. S. Baldwin, A. Reynaud, and R. F. Hess. The experiment was conducted by A. E. Finn. The analysis was performed by A. S. Baldwin. The manuscript was drafted by A. E. Finn, A. S. Baldwin, and R. F. Hess. All authors revised the manuscript.

